# The age-related susceptibility to postoperative delirium quantified by bispectral electroencephalography (BSEEG) correlates with postoperative delirium-like behavior in mice

**DOI:** 10.1101/2025.03.10.642417

**Authors:** Tsuyoshi Nishiguchi, Kyosuke Yamanishi, Akiyoshi Shimura, Tomoteru Seki, Takaya Ishii, Bun Aoyama, Nipun Gorantla, Hieu Dinh Nguyen, Nathan James Phuong, Takehiko Yamanashi, Massaaki Iwata, Gen Shinozaki

**Affiliations:** Department of Psychiatry and Behavioral Sciences, Stanford University School of Medicine, Palo Alto, CA, USA; Department of Neuropsychiatry, Faculty of Medicine, Tottori University, Yonago, Tottori, Japan; Department of Neuropsychiatry, School of Medicine, Hyogo Medical University, Nishinomiya, Hyogo, Japan; Department of Psychiatry, Tokyo Medical University, Shinjuku-ku, Tokyo, Japan; iPS Cell-Based Drug Discovery Group, Regenerative & Cellular Medicine Kobe Center, Sumitomo Pharma Co., Ltd., Osaka, Osaka, Japan; Division of Anesthesiology, National Hospital Organization Kochi Hospital, Kochi, Kochi, Japan; Department of Anesthesiology and Intensive Care Medicine, Kochi Medical School, Nankoku, Kochi, Japan

**Author notes:** **Corresponding author:** Name: Gen Shinozaki, M.D. Mailing Address: 3165 Porter Drive, Palo Alto, CA, 94304, USA.

**Keywords:** Delirium, Postoperative delirium (POD), Electroencephalography (EEG), Bispectral electroencephalography (BSEEG), Behavioral test, Mice

## Abstract

Postoperative delirium (POD) is a severe neuropsychiatric state characterized by acute fluctuating various mental symptoms. Despite its prevalence, the underlying mechanisms of POD remain largely unknown, and effective biomarkers for its detection are lacking. We have successfully developed a novel delirium detection method, the bispectral electroencephalography (BSEEG) method, which has shown excellent performance in delirium detection and outcome prediction. From there, similar to the clinical situation, we hypothesize that BSEEG scores can serve as indicators of POD-like states in mice. This study investigated the correlation between POD-like behavior and BSEEG score in a mouse model following EEG head mount implantation surgery.

2-3 and 22-23 month-old male C57BL/6J mice underwent EEG head-mount implantation surgery, followed by EEG monitoring and a battery of behavioral tests, including Buried Food Test, Open Field Test, and Y-maze, to assess POD-like behavior. We measured BSEEG scores and analyzed the correlation between these scores and the behavior measurements.

The results showed that BSEEG scores correlated with attention deficit and decreases in locomotor activity in young mice, whilst BSEEG scores only correlated with a decrease in locomotor activity in aged mice. Notably, composite Z scores representing delirium severity also showed a correlation with BSEEG scores in young mice.

Our findings indicate that the BSEEG method can be an indicator of POD-like states consistent with those measured by behavior tests. This study provides a novel preclinical framework for understanding the pathophysiology of POD and underscores the potential of BSEEG as a valuable tool for delirium detection and severity assessment.

## 1. Introduction

Delirium is a severe neuropsychiatric syndrome characterized by the acute onset of deficits in attention, memory, and cognition(Wilson et al., 2020). It correlates with adverse events, including elevated mortality, increased likelihood of post-discharge institutionalization, and persistent cognitive decline, potentially leading to dementia(Witlox et al., 2010). Particularly in older adults, postoperative delirium (POD) presents as a prevalent post-surgical complication(American Geriatrics Society Expert Panel on Postoperative Delirium in Older Adults, 2015). Despite its profound effects, the pathophysiological mechanisms underlying POD remain elusive(Safavynia et al., 2018). Diagnosis of POD predominantly relies on clinical assessments and expert judgment due to a lack of reliable biomarkers(Setters and Solberg, 2017). Approximately half of all delirium cases remain undiagnosed(Spronk et al., 2009), underscoring a critical need for improved detection and severity quantification methods for delirium.

Our study has introduced an innovative approach through the application of a novel bispectral electroencephalography (BSEEG) method designed to detect delirium and predict patient outcomes by analyzing characteristic features of delirium based on the ratio of 3Hz power to 10Hz power to capture slow brainwave activity(Shinozaki et al., 2018). This BSEEG method, which utilizes a simple one-channel EEG, has demonstrated its effectiveness in delirium detection and is strongly linked with key clinical outcomes, such as mortality, hospital length of stay, and post-discharge disposition(Shinozaki et al., 2019a; Yamanashi et al., 2021a). Validated across over 1,000 patients, the BSEEG method has shown promising application to clinical practice(Nishizawa et al., 2023).

Building on this success in clinical application, we have expanded our investigation into preclinical studies to further delineate the pathophysiology of delirium, aiming to facilitate the development of targeted preventative and therapeutic interventions. Some studies have revealed that surgical procedures or the administration of lipopolysaccharide (LPS) trigger systemic and neuro-inflammatory responses, inducing a state akin to delirium(Field et al., 2012; Yang et al., 2020). Our previous study demonstrated that the BSEEG methodology applied in a mouse model captures an increase in the BSEEG score following LPS administration, consistent with delirium in patients(Nishiguchi et al., 2024c; Yamanashi et al., 2021c). Additionally, we have reported that the BSEEG method following EEG head mount implantation surgery in mice exhibits a prolonged increase in the BSEEG score over several days postoperatively, followed by a gradual recovery in the BSEEG score over a few days to several days depending on their age(Nishiguchi et al., 2024a). To further demonstrate the validity and usefulness of the BSEEG method in a mouse model, our group also showed a correlation between the BSEEG score and LPS induced delirium-like behavior based on behavior tests(Nishiguchi et al., 2024b).

However, the correlation between the BSEEG score and POD-like behavior after the EEG head mount implantation surgery model has never been investigated. We hypothesized that the BSEEG score would correlate with POD-like behavior and that the BSEEG method could be a reliable indicator of a POD-like state in the surgery mouse model. This study aims to clarify the relationship between the BSEEG score and POD-like behavior in this surgical model, offering valuable insights into the broader applicability of the BSEEG method in the preclinical investigation of delirium.

## 2. Methods and materials

### 2.1. Animals and housing

Young male C57BL/6J mice (2–3 months old (n = 15)) were purchased from Jackson Laboratory, and aged male C57BL/6J mice (22-23 months old (n=10) were delivered from the National Institute on Aging. They were housed at Stanford University’s animal facility, and 4-5 mice shared a cage with free access to food and water before EEG head-mount implantation surgery. Post-surgery, each was housed individually. The environment was maintained in a 12:12 h light-dark cycle, a temperature of 20-26, and a humidity of 30-60%.

### 2.2. Experimental procedure

#### 2.2.1 The schedule of the experiment

The EEG head-mount implantation surgery was performed during the daytime. EEG recording began at night on the surgery day, spanning postoperative day (PO-Day) 0 to 8, with PO-Day 0 as the surgery day. The battery of the behavioral tests was performed on PO-Day 1, 3, 5, 7 (Fig. 1a).

**Fig. 1.**
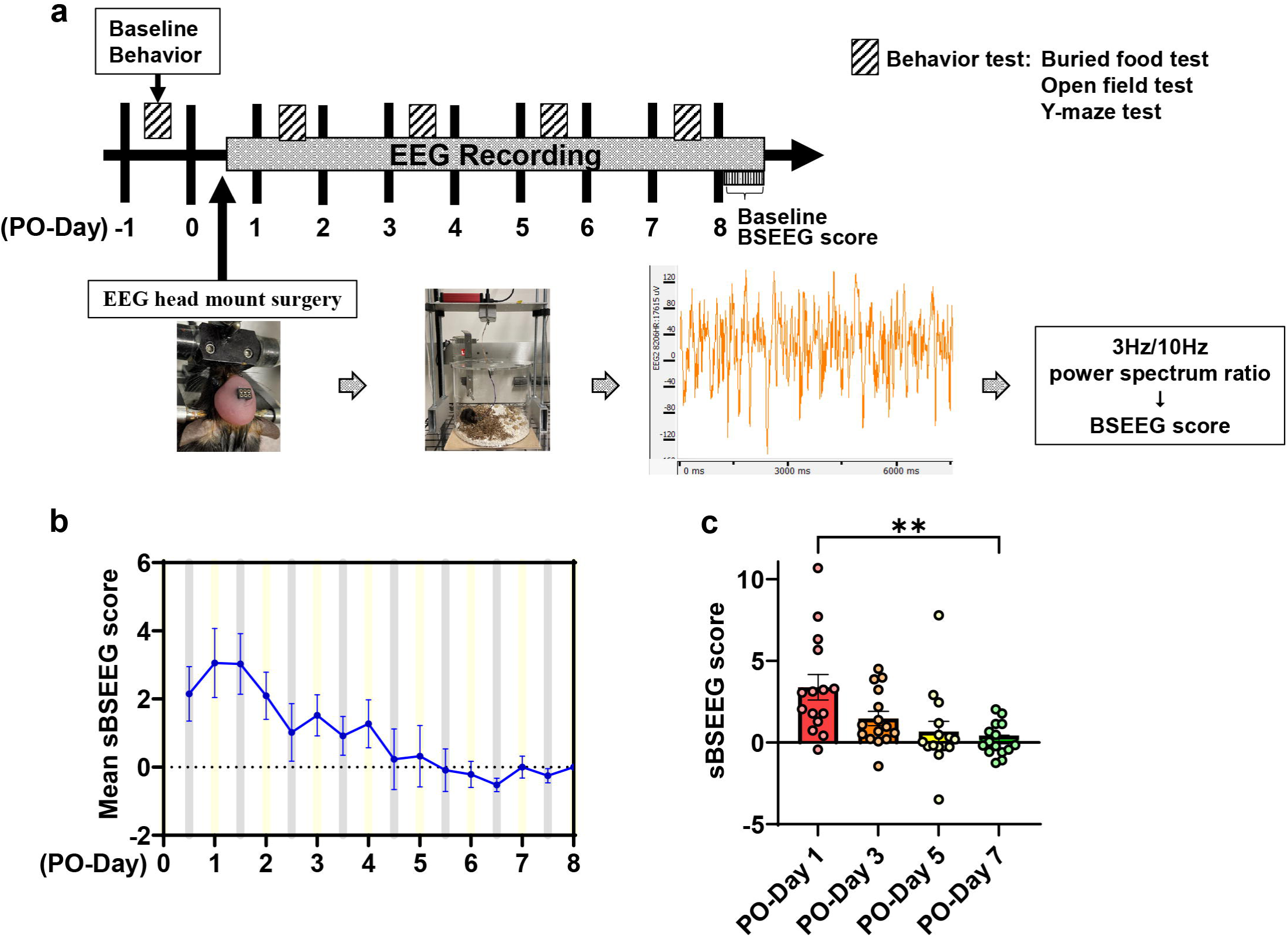
Schematic diagram of the experiment and the postoperative sBSEEG score in young mice. (a) Schematic diagram of the experiment and the postoperative sBSEEG score. (b) The mean postoperative sBSEEG score time course in young mice. (c) The differences in sBSEEG scores comparing PO-Day 1, PO-Day 3, PO-Day 5, and PO-Day 7 groups in young mice(mean = 3.38 (PO-Day 1), 1.47 (PO-Day 3), 0.67 (PO-Day 5), 0.19 (PO-Day 7); p = 0.0012 (PO-Day 1 vs. PO-Day 7))

#### 2.2.2. EEG electrode head mount placement surgery

EEG head mount implantation surgery was conducted as described in the previous studies(Buchanan et al., 2014; Nishiguchi et al., 2024a; Nishiguchi et al., 2024b; Nishiguchi et al., 2024c; Purnell et al., 2017; Yamanashi et al., 2021c) (see the Supplemantal Methods for details).

#### 2.2.3. EEG signal processing for BSEEG scores

Raw EEG was digitally converted into power spectral density to calculate BSEEG scores. We used the Sirenia^®^ software to record and export raw data into a European Data Format (EDF) file. Similar to a previous report of a web-based delirium detection application for human(Shimura et al., 2024), we processed an EDF file through our web-based tool (https://sleephr.jp/gen/mice7days/) for mice to calculate BSEEG scores. Our web-based tool is an automatic calculator for the BSEEG score based on the ratio of 3 Hz to 10 Hz power(Nishiguchi et al., 2024a).

#### 2.2.4. BSEEG score

In this study, we defined a BSEEG score as the average BSEEG score for every 12 hours from 7 a.m. to 7 p.m. to highlight the regular diurnal changes seen along with the circadian rhythm reported in the previous study(Yamanashi et al., 2021c).

#### 2.2.5. Standardized BSEEG (sBSEEG) score

To highlight the differences in the BSEEG scores compared to the steady state achieved post-surgery, we defined the standardized BSEEG (sBSEEG) score as the difference between daytime/night BSEEG scores on each PO-Day and daytime BSEEG scores on PO-Day 8. This means that the daytime BSEEG score of each mouse on PO-Day 8 was defined as sBSEEG score = 0. We adopted this approach because the baseline EEG cannot be recorded before head mount surgery and because most postoperative BSEEG scores recovered enough to reach steady in around one week after surgery in our previous study(Nishiguchi et al., 2024a). Therefore, most mice were estimated to fully recover and reach a steady state with their BSEEG scores by PO-Day 8.

### 2.3. Behavioral tests

The battery of behavioral tests was performed as described in a previous study(Nishiguchi et al., 2024b). Each test was performed 1 day before and 1, 3, 5, and 7 days after the surgery. The evaluation items of each behavioral test were calculated as a relative percentage of the baseline. The correlation between behavioral test endpoints and BSEEG score changes after the surgery was examined. The movement parameters of the mouse were counted by experimenters and the SMART 3.0 tracking system software (Panlab Harvard Apparatus, Holliston, MA, USA).

#### 2.3.1. Buried Food Test

BFT was introduced as a module to specifically evaluate attention in animals, which is typically disturbed as a result of delirium. Although it was originally used to test olfaction of mice(Machado et al., 2018; Yang and Crawley, 2009), many studies in the field have widely used this method to quantify attention and organized thinking(Lu et al., 2020; Matsumoto et al., 2021; Zhang et al., 2019; Zhang et al., 2018). This test was performed as described in a previous study(Nishiguchi et al., 2024b) (see the Supplemantal Methods for details).

#### 2.3.2. Open Field Test

The open field test was performed as described in a previous study(Nishiguchi et al., 2024b), and assessed the mice’s locomotor activity, motor function, and natural behavior(Combrinck et al., 2002; Cunningham et al., 2009; Peng et al., 2016) (see the Supplemantal Methods for details).

#### 2.3.3. Y maze test

The Y maze test is a test meant to quantify spatial and working memory based on spontaneous behavior, with the advantage that it does not require any reward or punishment. It can be performed quickly with a single learning phase(Wahl et al., 2017). The Y maze test was performed as described in a previous study(Nishiguchi et al., 2024b) (see the Supplemantal Methods for details).

#### 2.3.4. Composite Z score

The battery of the behavioral tests was reported as postoperative impairments in mice consistent with the Confusion Assessment Method (CAM) feature in delirium. The composite Z score in the battery of the behavioral tests was also reported to be able to quantitatively describe the behaviors analogous to a delirium severity measurement(Peng et al., 2016). Each Z score of the behavioral parameter was calculated as Z = [Δ*X _PO-Day_*-MEAN (Δ*X _PO-Day 7_)*]/SD (Δ*X _PO-Day 7_)(Liu et al., 2022)*. In the formula, Δ*X _PO-Day 7_* was the change score that the score of mice in the PO-Day7 group minus the score at the baseline; Δ*X _PO-Day_* was the change score that the score of mice in PO-Day 1, 3, 5 groups minus the score at baseline; MEAN (Δ*X _PO-Day 7_)* was the mean of Δ*X _PO-Day 7_*; and SD (Δ*X _PO-Day 7_)* was the standard deviation of Δ*X _PO-Day 7_*. The composite Z score for the mouse was calculated as the sum of the values of 6 Z scores (latency to pellet, time spent in center, latency to center, freezing time, entries in novel arm, and duration in novel arm) normalized with the SD for that sum in the saline. Given that the reduction in time spent in the center, the freezing time (OFT), the duration, and the entries in novel arm (Y maze) indicate impairment of the behavior, we multiplied the Z score values representing these behaviors by −1 before calculating the composite Z score using these values.

### 2.4. Statistical analysis

Statistical analysis was performed using GraphPad Prism version 10. Error bars were shown as means ± standard error of the means (SEM). Normality was assessed using the Shapiro-Wilk normality test. The differences between groups were analyzed, as PO-Day7 was defined as control, with either one-way repeated measures ANOVA using the Geisser-Greenhouse correction for possible violations of sphericity followed by Dunnett’s multiple comparison tests, where data were normally distributed, or with the Friedman test followed by Dunn’s multiple comparison tests, where data were not normally distributed. A P value less than 0.05 was marked with an asterisk and considered statistically significant, and P values are indicated as follows in figures: *P < 0.05, **P < 0.01, ***P < 0.001. Correlations were calculated using Spearman’s test. Mice that died post-surgery were excluded from the analysis.

## 3. Results

### 3.1. BSEEG score in young mice

Mean postoperative sBSEEG scores time course showed its peak on PO-Day 0 to 1. After PO-Day 1, mean postoperative sBSEEG scores gradually recovered (Fig. 1b). Postoperative sBSEEG scores were significantly higher on PO-Day 1 than on PO-Day 7 (Fig. 1c). Many mice showed the peak of raw BSEEG scores within a few days after surgery (Fig. S1a). Mean postoperative raw BSEEG scores time course also showed its peak on PO-Day 0 to 1. After PO-Day 1, the average of the postoperative raw BSEEG scores gradually decreased (Fig. S1b). Postoperative raw BSEEG scores also were significantly higher on PO-Day 1 than on PO-Day 7 (Fig. S1c).

### 3.2. Behavioral tests in young mice

Latency to pellet in BFT was significantly higher on PO-Day 1 than on PO-Day 7, and on PO-Day 3 than on PO-Day 7 (Fig. 2a). Total distance in OFT was significantly lower on PO-Day 1 than on PO-Day 7 and on PO-Day 3 than on PO-Day 7 (Fig. 2b). There were no significant differences on each day in time spent in center (Fig. 2c), freezing time (Fig. 2d), latency to center (Fig. 2e) in OFT, duration in novel arm (Fig. 2f), and entries in novel arm (Fig. 2g) in Y maze.

**Fig. 2.**
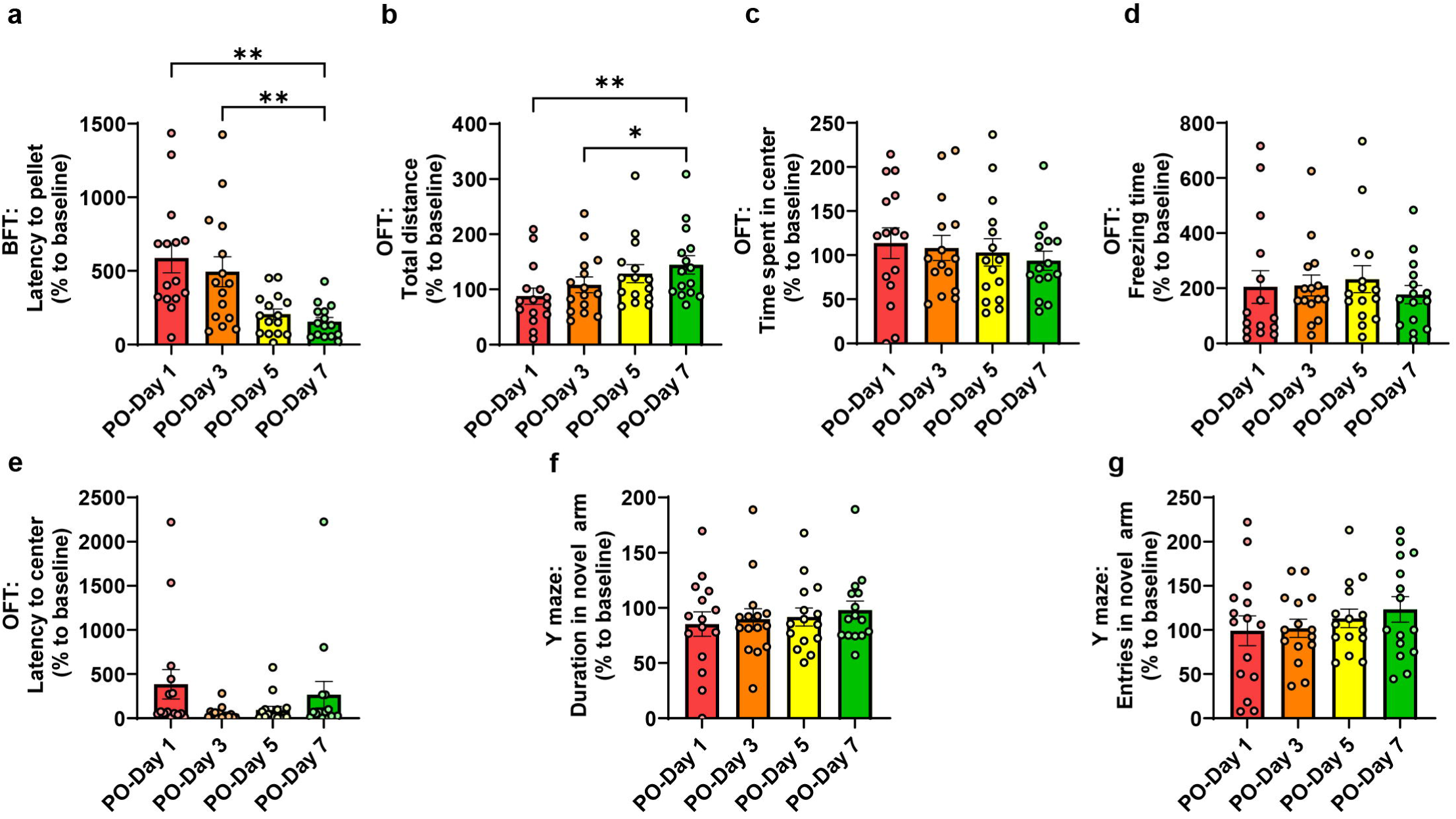
The differences in behavioral tests comparing PO-Day 1, PO-Day 3, PO-Day 5, and PO-Day 7 groups in young mice. (a) Latency to pellet in BFT (mean = 586.5 % (PO-Day 1), 494.1 % (PO-Day 3), 206.5 % (PO-Day 5), 155.2 % (PO-Day 7); p = 0.0028 (PO-Day 1 vs. PO-Day 7), p = 0.01 (PO-Day 3 vs. PO-Day 7)). (b) Total distance in OFT (mean = 87.76 % (PO-Day 1), 108.0 % (PO-Day 3), 128.6 % (PO-Day 5), 144.5 % (PO-Day 7); p = 0.0021 (PO-Day 1 vs. PO-Day 7), p = 0.022 (PO-Day 3 vs. PO-Day 7)). (c) Time spent in center in OFT(mean = 113.5 % (PO-Day 1), 108.1 % (PO-Day 3), 103.1 % (PO-Day 5), 93.62 % (PO-Day 7)). (d) Freezing time in OFT (mean = 204.9 (PO-Day 1), 209.8 % (PO-Day 3), 232.6 % (PO-Day 5), 176.8 % (PO-Day 7)). (e) Latency to center in OFT (mean = 384.3 % (PO-Day 1), 52.02 % (PO-Day 3), 93.60 % (PO-Day 5), 265.9 % (PO-Day 7)). (f) Duration in novel arm in Y maze(mean = 85.32 % (PO-Day 1), 89.68 % (PO-Day 3), 91.75 % (PO-Day 5), 97.95 % (PO-Day 7)). (g) Entries in novel arm in Y maze (mean = 99.07 % (PO-Day 1), 101.8 % (PO-Day 3), 113.1 % (PO-Day 5), 123.3 % (PO-Day 7))

### 3.3. Correlations between BSEEG score and behavioral test in young mice

Next, we analyzed correlations between the sBSEEG score and behavioral tests. Latency to pellet in BFT showed a significant moderate positive correlation to sBSEEG score (Fig. 3a). Also, total distance in OFT showed a significant moderate negative correlation to the sBSEEG score (Fig. 3b). There was no correlation between sBSEEG score and time spent in center, freezing time and latency to center in OFT or duration and entries in novel arm in Y maze (Fig. 3c, 3d, 3e, 3f, and 3g).

**Fig. 3.**
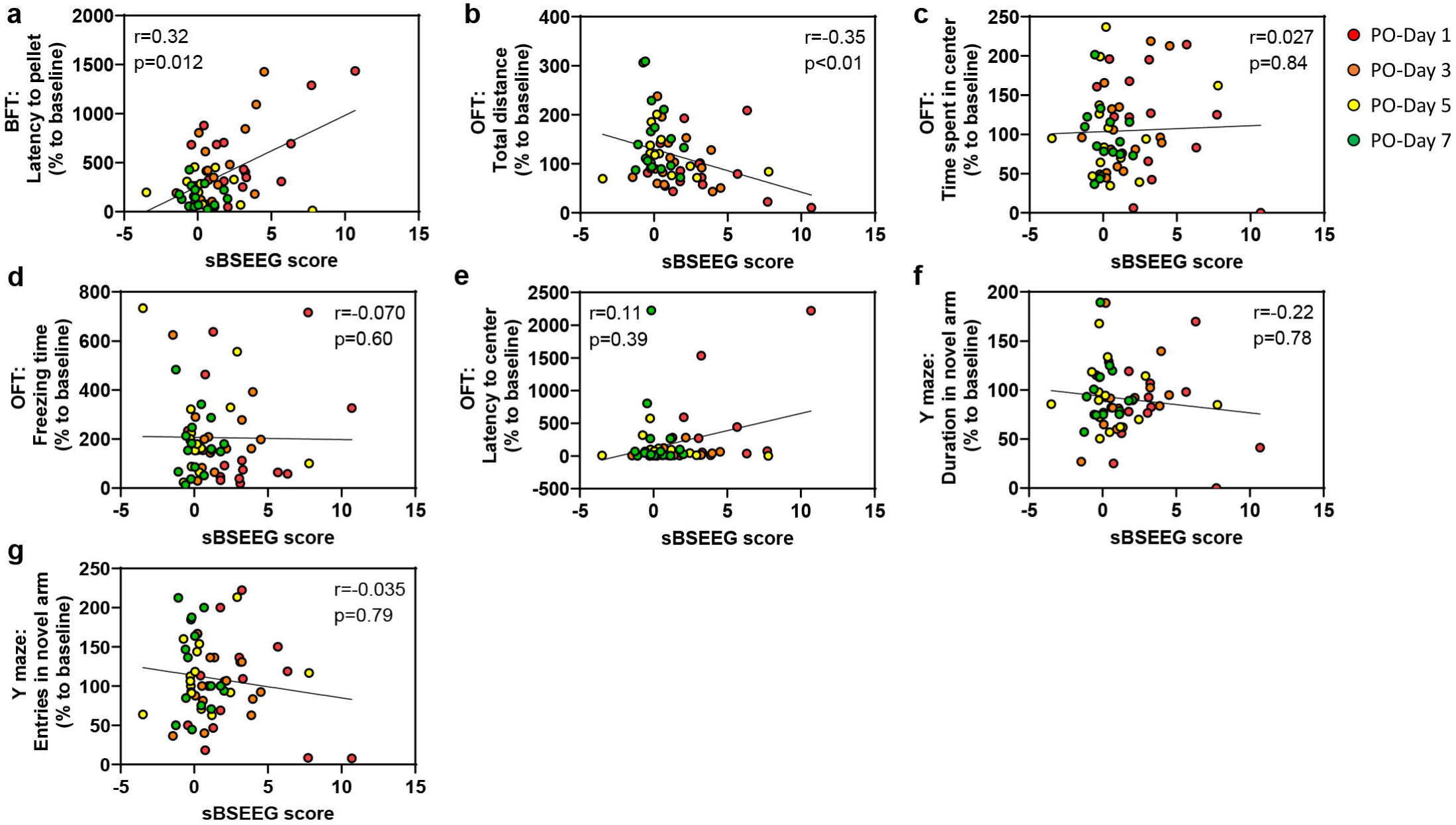
Correlations between sBSEEG score and behavioral tests in young mice. Correlations between sBSEEG scores and (a) latency to pellet in BFT, (b) total distance in OFT, (c) time spent in center in OFT, (d) freezing time in OFT, (e) latency to center in OFT, (f) duration in novel arm in Y maze, and (g) entries in novel arm in Y maze

Finally, we calculated the composite Z score and analyzed the correlation between the sBSEEG score and the composite Z score. The composite Z score was significantly higher on PO-Day 1 than on PO-Day 7 (Fig. 4a). The Composite Z score showed a significant moderate positive correlation to the sBSEEG score and raw BSEEG score (Fig. 4b)(Fig. S2).

**Fig. 4.**
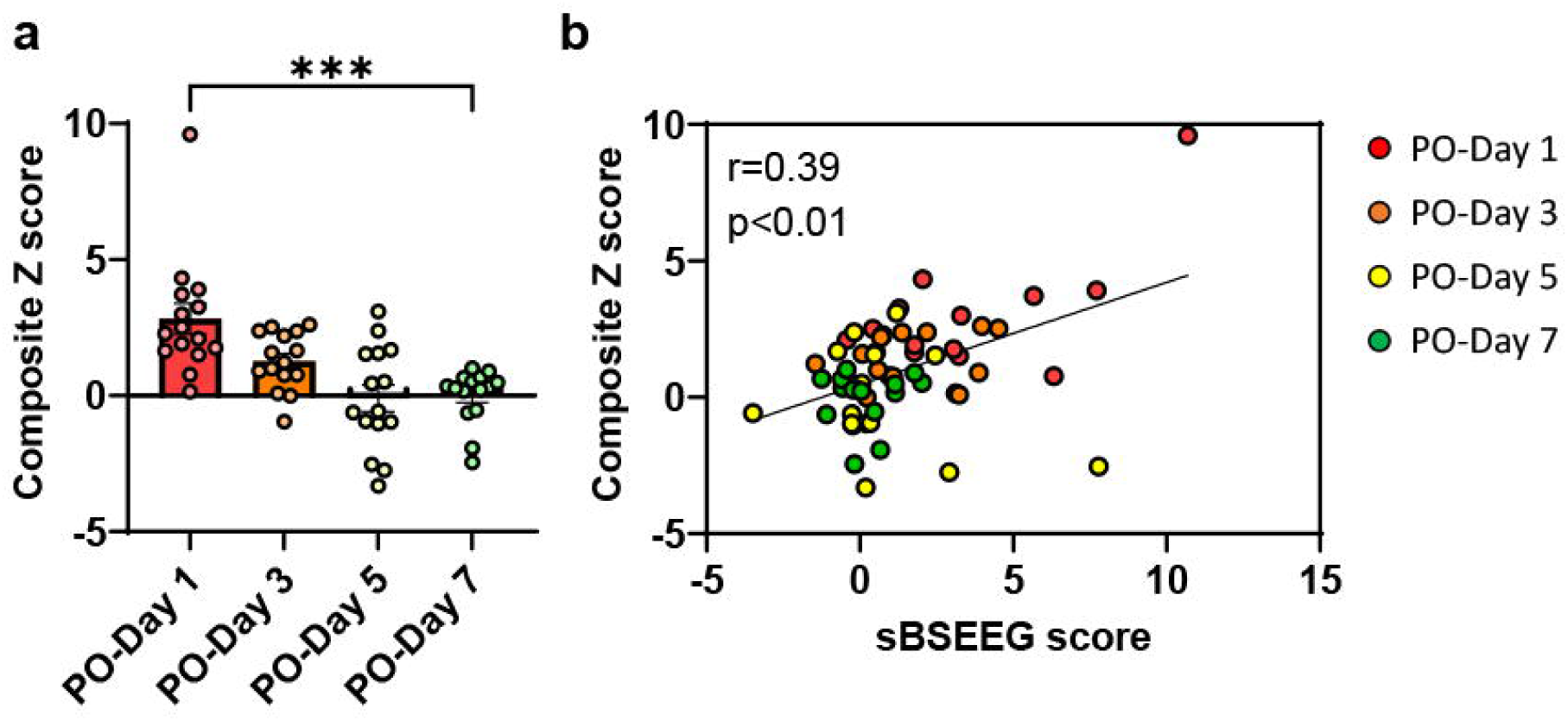
Composite Z score in young mice. (a) The differences in composite Z score comparing PO-Day 1, PO-Day 3, PO-Day 5, and PO-Day 7 groups (mean = 2.83 (PO-Day 1), 1.27 (PO-Day 3), −0.10 (PO-Day 5), 0.00 (PO-Day 7); p = 0.0001 (PO-Day 1 vs. PO-Day 7)). (b) Correlation between sBSEEG scores and composite Z scores

### 3.4. BSEEG score in aged mice

Mean postoperative sBSEEG scores time course showed its peak on PO-Day 0 to 1, and mean postoperative sBSEEG scores gradually recovered (Fig.5a). Postoperative sBSEEG scores were significantly higher on PO-Day 1 than on PO-Day 7 and higher on PO-Day 3 than on Day 7 (Fig. 5b). Many aged mice showed the peak of raw BSEEG scores within a few days after surgery as young mice did (Fig. S3a). However, mean postoperative raw BSEEG scores in aged mice were higher than those in young mice (Fig. S3b). Postoperative raw BSEEG scores were also significantly higher on PO-Day 1 than on PO-Day 7 and on PO-Day 3 than on PO-Day7 (Fig. S3c).

**Fig. 5.**
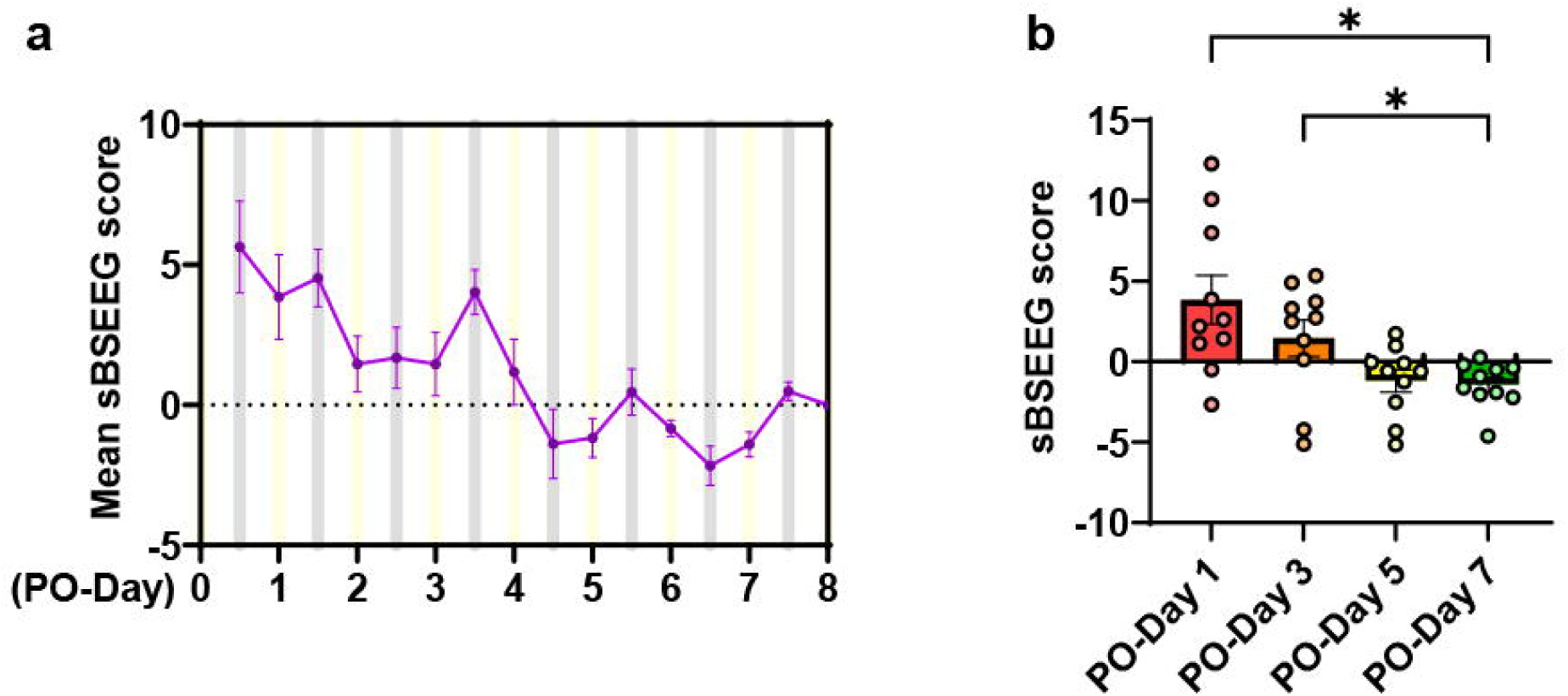
The postoperative sBSEEG score in aged mice. (a) The time course of mean postoperative sBSEEG score in aged mice. (b) The differences in sBSEEG scores comparing PO-Day 1, PO-Day 3, PO-Day 5, and PO-Day 7 groups in aged mice (mean = 3.85 (PO-Day 1), 1.47 (PO-Day 3), −1.18 (PO-Day 5), −1.41 (PO-Day 7); p = 0.023 (PO-Day 1 vs. PO-Day 7), p = 0.021 (PO-Day 3 vs. PO-Day 7)).

### 3.5. Behavioral Tests in aged mice

There were no significant differences on each day in latency to pellet in BFT (Fig. 6a). Total distance in OFT was significantly lower on PO-Day 1 than on PO-Day 7, and on PO-Day 3 than on PO-Day 7 (Fig. 6b). There were no significant differences on each day in time spent in center (Fig. 6c). Freezing time in OFT was significantly higher on PO-Day 1 than on PO-Day 7, and on PO-Day 3 than on PO-Day 7 (Fig. 6d). There were no significant differences on each day in latency to center (Fig. 6e) in OFT, and in duration in novel arm (Fig. 6f) in Y maze. Entries in novel arm was significantly higher on PO-Day1 than on PO-Day 7 (Fig. 6g) in Y maze.

**Fig. 6.**
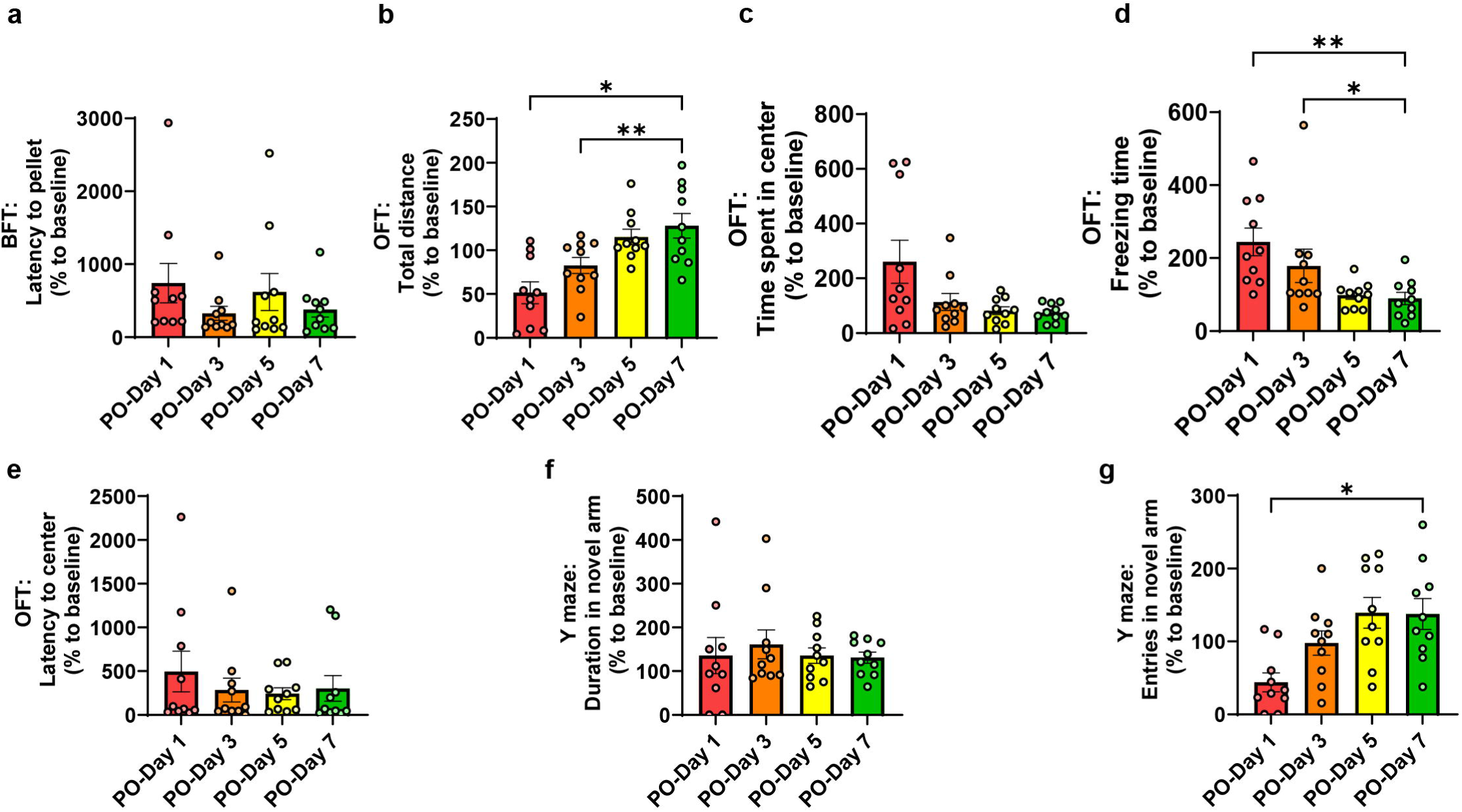
The differences in behavioral tests comparing PO-Day 1, PO-Day 3, PO-Day 5, and PO-Day 7 groups in aged mice. (a) Latency to pellet in BFT (mean = 741 % (PO-Day 1), 327 % (PO-Day 3), 619 % (PO-Day 5), 377 % (PO-Day 7)). (b) Total distance in OFT (mean = 51 % (PO-Day 1), 83 % (PO-Day 3), 115 % (PO-Day 5), 128 % (PO-Day 7); p = 0.018 (PO-Day 1 vs. PO-Day 7), p = 0.0027 (PO-Day 3 vs. PO-Day 7)). (c) Time spent in center in OFT (mean = 260 % (PO-Day 1), 114 % (PO-Day 3), 82 % (PO-Day 5), 75 % (PO-Day 7)). (d) Freezing time in OFT(mean = 244 % (PO-Day 1), 178 % (PO-Day 3), 98 % (PO-Day 5), 89 % (PO-Day 7); p = 0.0016 (PO-Day 1 vs. PO-Day 7), p = 0.017 (PO-Day 3 vs. PO-Day 7)). (e) Latency to center in OFT (mean = 496 % (PO-Day 1), 284 % (PO-Day 3), 242 % (PO-Day 5), 302 % (PO-Day 7)). (f) Duration in novel arm in Y maze (mean = 136 % (PO-Day 1), 161 % (PO-Day 3), 136 % (PO-Day 5), 131 % (PO-Day 7)). (g) Entries in novel arm in Y maze (mean = 44 % (PO-Day 1), 98 % (PO-Day 3), 139 % (PO-Day 5), 138 % (PO-Day 7), p = 0.012 (PO-Day 1 vs. PO-Day 7)).

### 3.6. Correlations between BSEEG score and behavioral test in aged mice

Latency to pellet in BFT did not show a correlation to the sBSEEG score (Fig. 7a). Total distance showed a significant moderate negative correlation to the sBSEEG score (Fig. 7b). There was no correlation between the sBSEEG score and time spent in center in OFT (Fig. 7c). Freezing time in OFT showed a significant moderate correlation to the sBSEEG score (Fig. 7d). There was no correlation between the sBSEEG score and Latency to center in OFT (Fig. 7e). There was no correlation between the sBSEEG score and duration in Y maze (Fig. 7f). Entries in novel arm in Y maze showed a significant moderate negative correlation to the sBSEEG score (Fig. 7g).

**Fig. 7.**
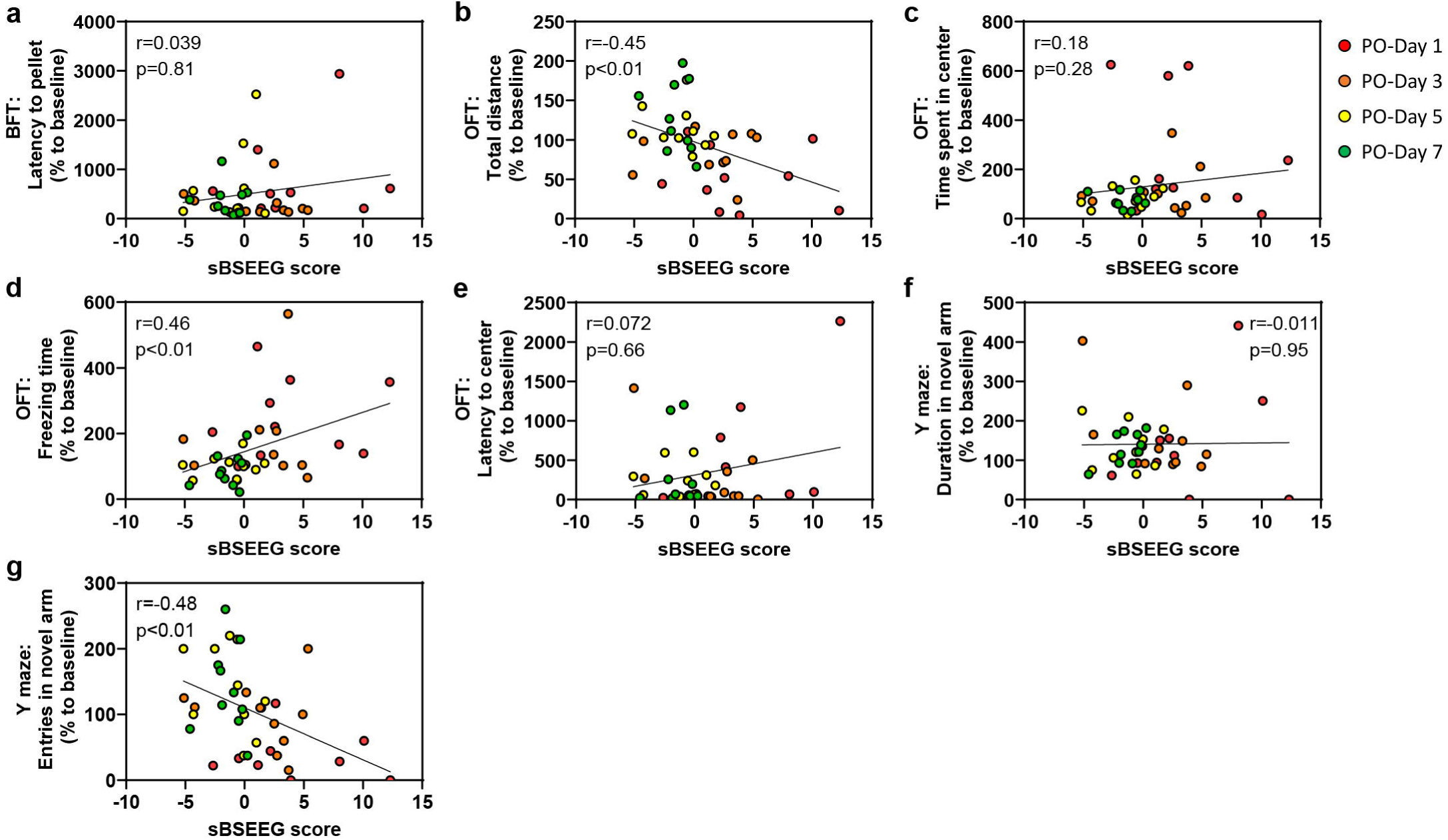
Correlations between sBSEEG score and behavioral tests in aged mice. Correlations between sBSEEG scores and (a) latency to pellet in BFT, (b) total distance in OFT, (c) time spent in center in OFT, (d) freezing time in OFT, (e) latency to center in OFT, (f) duration in novel arm in Y maze, and (g) entries in novel arm in Y maze.

We calculated the composite Z score and analyzed the correlation between the sBSEEG score and the composite Z score. There were no significant differences on each day in the composite Z score (Fig. 8a). The Composite Z score did not show a correlation to the sBSEEG score raw BSEEG score (Fig. 8b)(Fig. S4).

**Fig. 8.**
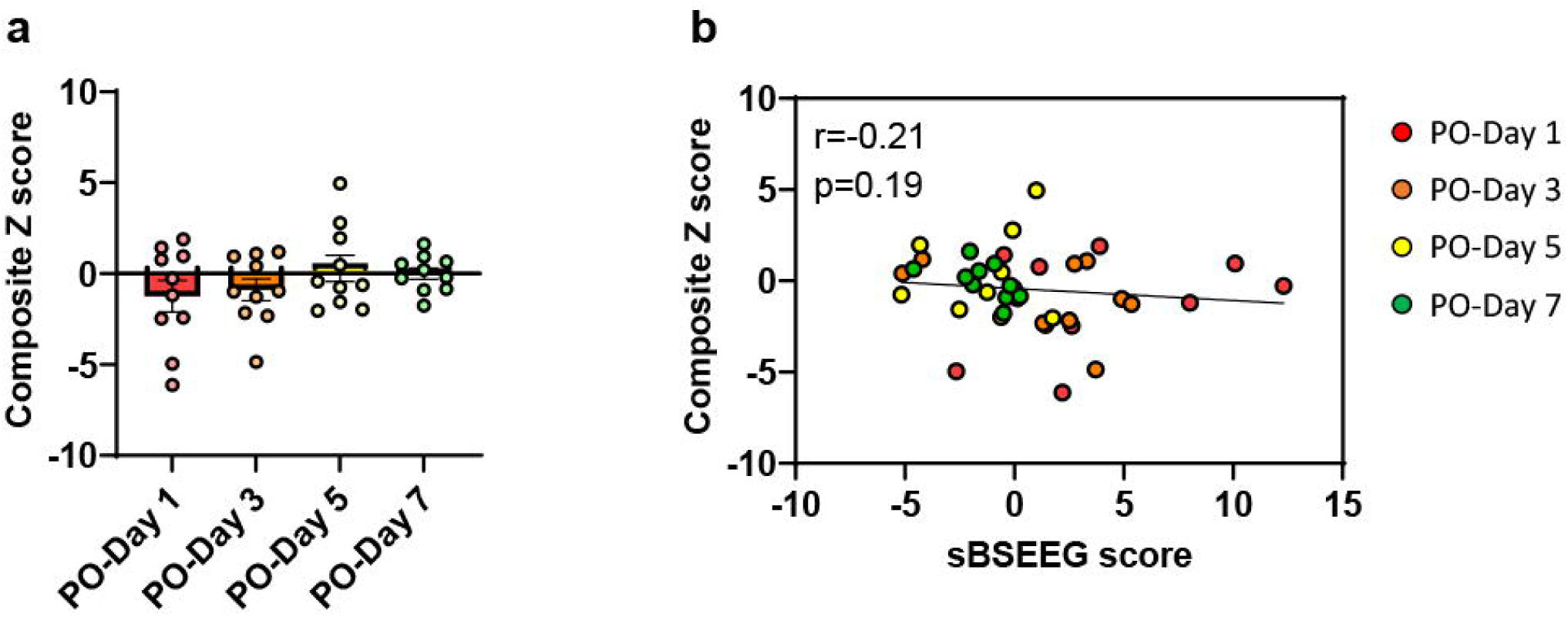
Composite Z score in aged mice. (a) The differences in composite Z score comparing PO-Day 1, PO-Day 3, PO-Day 5, and PO-Day 7 groups (mean = −1.24 (PO-Day 1), −0.89 (PO-Day 3), 0.28 (PO-Day 5), 0.00 (PO-Day 7)). (b) Correlation between sBSEEG scores and composite Z scores.

## 4. Discussion

This study is the first report to verify the level of correlation between postoperative BSEEG score and delirium-like behavior measured by several behavioral tests over the time course of recovery after surgical intervention, supporting our hypothesis that the BSEEG method can serve as an indicator of a postoperative delirium-like state after surgery in mice.

In the present study, the BSEEG scores are calculated from a ratio of approximately 3Hz to 10Hz, as conducted in our previous reports in both human studies and mouse studies(Nishiguchi et al., 2024a; Nishiguchi et al., 2024c; Shinozaki et al., 2019b; Shinozaki et al., 2018; Yamanashi et al., 2021b; Yamanashi et al., 2021c). A higher BSEEG score indicates a higher presence of slow wave components in EEG. This method is based on slow wave components in EEG being more prominent under the disturbance of consciousness observed in delirium states. Like our previous study, this study also showed that postoperative BSEEG scores increased for several days and decreased gradually over time after that period(Nishiguchi et al., 2024a).

In this postoperative behavioral study, to evaluate delirium-like states in mice, we followed the protocols commonly used in the literature and conducted a battery of three behavioral tests, instead of a single behavioral test. This is in line with CAM that evaluates a combination of symptoms typically observed in delirium in humans, including acute fluctuations in consciousness, impaired attention, and impaired thinking(Peng et al., 2016). Some previous studies reported that the battery of postoperative behavioral tests could capture significant differences in levels of delirium-like behavior, even in aged mice maintaining the level of their locomotor activity(Li et al., 2021; Zhou et al., 2020). Contrary to those reports, although we showed that the battery of postoperative behavioral tests could capture significant differences in levels of delirium-like behavior in young mice, we could not capture such differences in aged mice for their level of delirium-like behavior and could only capture their differences in locomotor activity. This difference in results between those studies and our study may be due to the differences in surgery types and the time points of behavioral tests.

In our study, among young mice, there were significant differences between PO-Day 1 and PO-Day 7 in the latency to pellet in BFT, which is an indicator of impaired attention. These results were also significantly correlated with the BSEEG scores. Also, there was a significantly negative correlation between the BSEEG scores and total distance in OFT, which is an indicator of locomotor activity. On the other hand, the duration in novel arm in Y maze, a measure of spatial cognitive impairment, showed a trend toward improvement from PO-Day 1 to PO-Day 7, but it did not reach significance. These results suggest that in the POD-like model after EEG head-mount surgery, attention impairment was prominent as a delirium-like symptom, and the BSEEG score strongly reflected the attention impairment. Furthermore, the composite Z score that was an indicator of delirium severity significantly decreased from PO-Day 1 to PO-Day 7, and showed a positive correlation with the BSEEG score. Thus, these results suggest that the BSEEG score has the potential to be an effective indicator of POD-like behavior.

On the other hand, there were no obvious correlations between BSEEG scores and delirium-like behaviors in aged mice, especially latency to pellet in BFT and composite Z score, which are thought to be important measures in the delirium model, although there were good correlations in young mice. Instead, BSEEG scores in aged mice only negatively correlated with the tests of locomotor activity that were total distance and freezing time in OFT and entries in novel arm in Y maze. Therefore, locomotor activity changes could be one of the most reliable tests of delirium-like behavior. BSEEG scores might be also able to capture the extent of systemic physical burden that leads to decreases in locomotor activity while notable decreases in locomotor activity might have reduced the sensitivity of other behavioral tests in aged mice.

Decreased locomotor activity in a POD mouse model was not observed in one previous study based on a simple laparotomy postoperative model(Li et al., 2021). However, it was observed in another previous study based on cardiac surgery(Jia et al., 2024). It is thought that this difference may be due to the greater trauma and inflammatory response induced by cardiac surgery(Hovens et al., 2016; Jia et al., 2024). Decreased locomotor activity in the POD mouse model of this study may also be due to the result of the greater trauma and inflammatory response by EEG head mount implantation surgery in addition to increased vulnerability in aged mice. The results of this study suggest that BSEEG scores reflect the ability to accurately measure physiological changes in the brain even though strong invasion and inflammation result in decreased locomotor activity that interferes with accurate assessment in behavioral tests. In other words, this BSEEG method may prove to be effective in assessing delirium-like states in a vulnerable model such as aged mice or mice model for neurodegenerative disorders including dementia.

We acknowledge several limitations of this study. First, although we were able to detect POD-like behavior in young mice, we were not able to detect it in aged mice except for a decrease in locomotor activity. Because delirium more commonly presents in older patients, we anticipated that aged mice would show delirium-like behavior more obviously. However, the behavioral tests in this study failed to capture delirium-like symptoms. It is likely because aged mice were more vulnerable than young mice and tended to show a more pronounced decrease in postoperative locomotor activity than young mice, and the behavioral tests were influenced by the reduced locomotor activity. Thus, the correlation between the sBSEEG score and delirium-like symptoms may not have been detectable. Second, although PO-Day 8 was used as the reference date for sBSEEG score = 0 to align with younger mice because both age groups showed to reached recovery by PO-Day 8 in our previous study(Nishiguchi et al., 2024a). However, postoperative recovery may not have been sufficient for aged mice in this experiment. Therefore, the sBSEEG scores among aged mice might have been falsely suppressed as we might have used a BSEEG score higher than true stable score at a later time after PO-Day8. Third, the typical diurnal changes in BSEEG scores that regularly go up in the daytime and down in the night were not detected clearly, although they were slightly detected around PO-Day 7 when we assume mice were in steady states.

Mice are nocturnal, and their physiological sleep cycle differs from that of humans in that they sleep during the daytime. It is also known that slow wave components are predominantly detected during sleep. Our previous study reported that there were regular diurnal changes on BSEEG scores in POD mouse model(Nishiguchi et al., 2024a). Therefore, it is possible that conducting behavioral tests every other day during the daytime in this study might have broken the physiological sleep of mice, making them awake in the daytime, and affecting their circadian rhythms and BSEEG scores. Fourth, we use only male mice in this study. In general, gender differences have not been associated with delirium risk(Scholz et al., 2016). Some reports indicate that male sex is a risk factor associated with delirium(Wang et al., 2021; Wittmann et al., 2022), while others reported that female sex is a risk factor associated with delirium(Fann et al., 2002; Zhang et al., 2020). Thus, gender differences in delirium risk are still controversial as they were not consistent across settings(Ormseth et al., 2023). Thus, we used only male mice in this study because including female mice may complicate age comparisons due to gender differences with changing female hormone levels with age. Fifth, repeating the same behavioral tests could lead to habituation and learning effects. The latency to palettes in BFT gradually declined in young mice. It might be because of habituation. On the other hand, old mice could not attain a significant decline in the latency to palettes in BFT. It might be age-related susceptibility to cognitive dysfunctions due to mild operative neurotrauma if there were impairments of habituation and learning effect. However, we took measures for BFT by putting a pellet in a random position. Also, previous studies did not show any obvious habituation and learning effects in repeating the same behavioral tests even with relatively short intervals(Illendula et al., 2020; Peng et al., 2016).

In conclusion, to capture the POD-like state by adapting the BSEEG method to the postoperative mouse model, we investigated the relationship between postoperative behavior and BSEEG scores in the study of the POD-like state after the EEG head mount surgery in mice.

We were able to show the chronological changes of the BSEEG scores in both age groups, with its initial increase soon after surgery for a few days, followed by a gradual decrease and recovery. We were also able to capture more prominent POD-like behavior in young mice, especially impaired attention, measured by behavioral tests. The BSEEG scores and behavioral test measures for delirium showed a correlation in young mice. In aged mice, those behavioral tests were not able to capture a correlation between the BSEEG score and the behavioral test for delirium, suggesting that the BSEEG method may serve as a useful approach to detect POD-like state in mouse models.

## Ethical approval

The experiments followed a protocol approved by the Institutional Animal Care and Use Committee (IACUC), known as Stanford’s Administrative Panel on Laboratory Animal Care (APLAC), accredited by the Association for Assessment and Accreditation of Laboratory Animal Care (AAALAC).

## Acknowledgements

We appreciate the support from Stanford University School of Medicine Veterinary Service Center (VSC).

## Data availability

Data is provided within the manuscript or supplementary information files.

## Author contributions

GS conceptualized and coordinated the entire research study. TN designed and conducted experiments, acquired data, analyzed data, and wrote the initial draft of the manuscript. AS programmed the web-based BSEEG score calculator. NG conducted experiments and analyzed data. KY, AS, TS, TI, BA, NG, HDN, TY, and MI critically reviewed the manuscript. TN and GS wrote the final version of the manuscript.

