## Supplemental Material for "The age-related susceptibility to postoperative delirium quantified by bispectral electroencephalography (BSEEG) correlates with postoperative delirium-like behavior in mice"

Nishiguchi et al.

**Supplementary Methods**

**EEG electrode head mount placement surgery**

The surgery was performed as described in a previous studies(Buchanan et al., 2014; Nishiguchi et al., 2024a; Nishiguchi et al., 2024b; Nishiguchi et al., 2024c; Purnell et al., 2017; Yamanashi et al., 2021). The mice were placed under isoflurane (1-3% inhalation) anesthesia, with the skull exposed. The head mount was placed on the midline, with its anterior holes at 2 mm anterior to bregma and its posterior holes at 2 mm anterior to lambda ± 2 mm. Four holes were bored into the skull with a 23-gauge needle through the head mount holes. Four screws were inserted into holes. Finally, the head mount was fixed with dental cement. Mice received postoperative analgesia with meloxicam 5 mg/kg by subcutaneous injection. Mice were connected to an EEG system after completion of surgery and anesthesia. EEG was recorded from the right frontal cortex using the right anterior screw (EEG2) and the right parietal cortex using the right posterior screw (EEG1). The left anterior screw at the left frontal cortex served as the ground, and the left posterior screw at the left parietal cortex served as the reference. We used signals from EEG2 because EEG2 was documented to be more sensitive than EEG1 in a previous study(Yamanashi et al., 2021).

**Buried Food Test (BFT)**

BFT was performed as described in a previous study(Nishiguchi et al., 2024b). 24 hours before the test, we removed all chow pellets from the home cage but did not remove the water bottle to provide water ad libitum. The test cage was prepared with clean bedding (3 centimeters high). We buried one pellet 0.5 centimeters below the surface of the bedding so that it was not visible. We used about 2 g of pellets of the same chow the animals were regularly fed with. The location of the food pellet was randomly changed every time a new mouse was introduced. We placed the mouse in the center of the test cage and measured the latency time. Latency time was defined as the duration from the time when the mouse was placed in the test cage until the mouse uncovered the food pellet and grasped it in the forepaws and teeth. Mice were allowed to consume the pellet they found and were then returned to their home cage. The maximum observation time was set at 10 minutes. If the mouse could not find the pellet within 10 minutes, the testing session ended, and the latency was defined as 600 seconds for that mouse. After each test, we exchange the test cage with a new one to prevent the transmission of olfactory cues. We also changed gloves after each test.

**Open Field Test (OFT)**

OFT was performed as described in a previous study(Nishiguchi et al., 2024b). The mouse was gently placed in the center of an open field (40 × 40 × 40 cm (width × length × height), Stoelting, Wood Dale, IL, USA) and allowed to move freely for 5 min. We analyzed the total distance moved (centimeters) as the indicator of locomotor activity, the time spent in center of the open field (seconds) as the indicator of anxiety and exploratory behavior, the freezing time (seconds) as the indicator of anxiety, exploratory behavior, and locomotor activity, and the first attempt latency to center of the open field (seconds) as the indicator of anxiety and exploratory behaviors. The floor of the open field was cleaned with 70% ethanol solution between each test.

**Y maze test**

The Y maze test was performed as described in a previous study(Nishiguchi et al., 2024b). The Y maze consisted of three arms, with an angle of 120 degrees between each arm (5 × 35 × 10 cm, Panlab Harvard Apparatus, Holliston, MA). The Y maze test consisted of 2 trials separated by an inter-trial interval (ITI). The first trial (training) was 10 minutes in duration, which allowed the mouse to explore 2 arms (the start arm and the other arm) of the maze, with the novel arm being blocked. After a 2-hour ITI, the second trial (retention) was conducted. For the second trial, the mouse was placed back in the maze in the same start arm with free access to all 3 arms for 5 minutes. The time spent in novel arm indicated the spatial recognition memory. The entries into the novel arm indicated spatial recognition memory and locomotor activity. Each of the arms of the Y maze was cleaned with 70% ethanol solution between trials.

**Supplementary Figures**

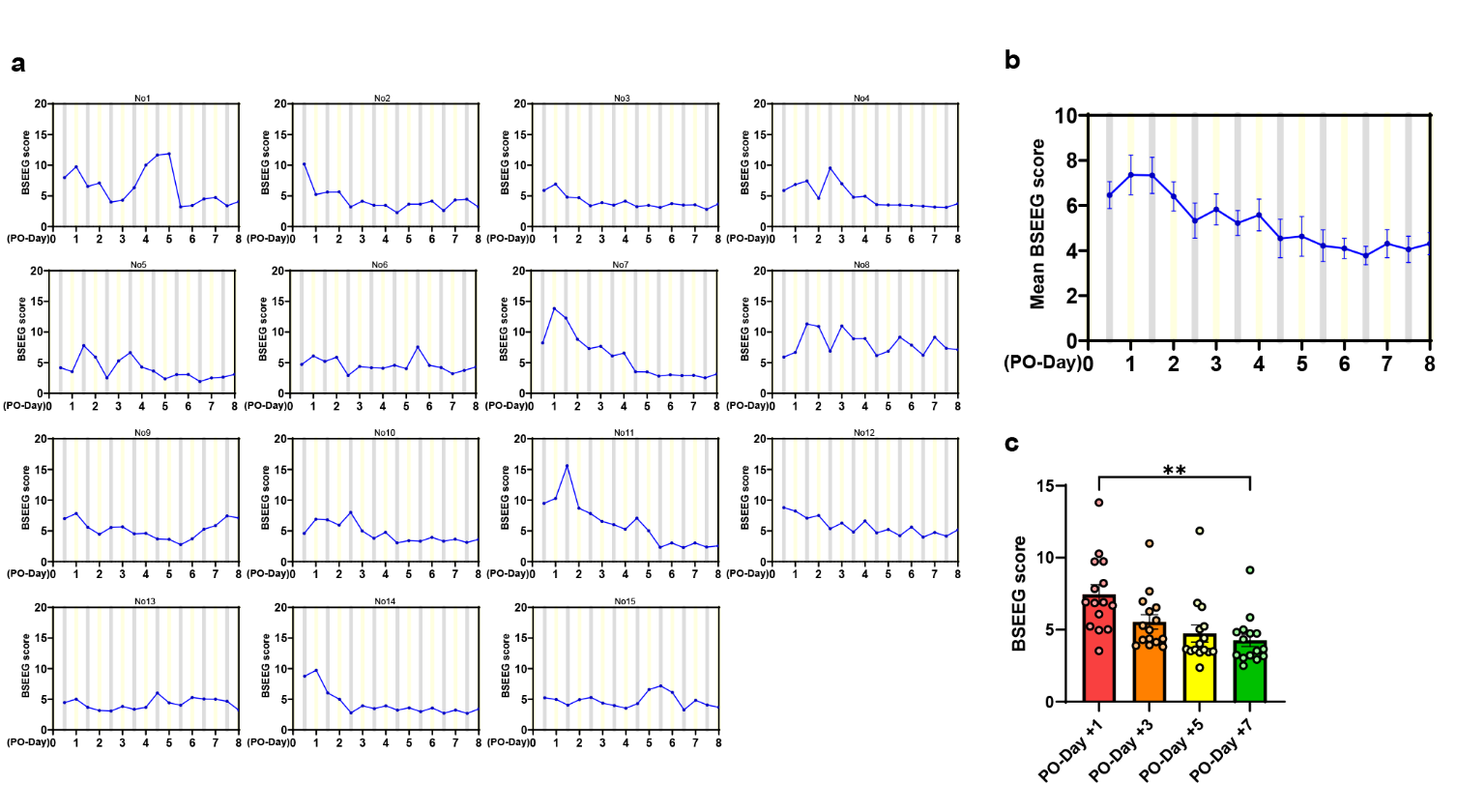

**Fig. S1** The postoperative raw BSEEG score in young mice. (a) The time courses of postoperative raw BSEEG score in each young mouse. Many mice showed the peak of raw BSEEG scores within a few days after surgery (No 2, 3, 4, 5, 7, 9, 10, 11, 12, 14). (b) The time course of mean postoperative raw BSEEG score in young mice. (c) The differences in raw BSEEG scores comparing PO-Day 1, PO-Day 3, PO-Day 5, and PO-Day 7 groups in young mice (mean = 7.46 (PO-Day 1), 5.55 (PO-Day 3), 4.75 (PO-Day 5), 4.26 (PO-Day 7); p = 0.0012 (PO-Day 1 vs. PO-Day 7))

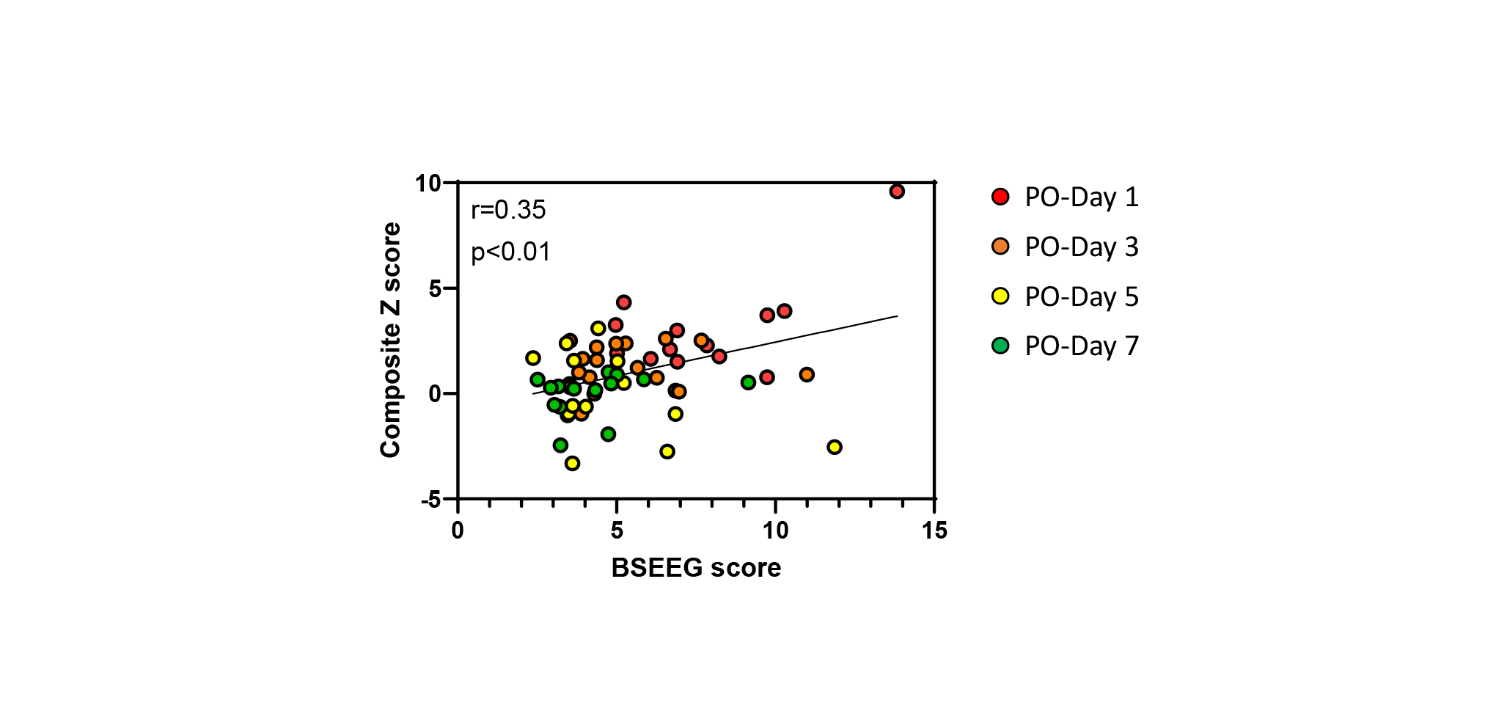

**Fig. S2** Correlation between BSEEG scores and composite Z scores in young mice

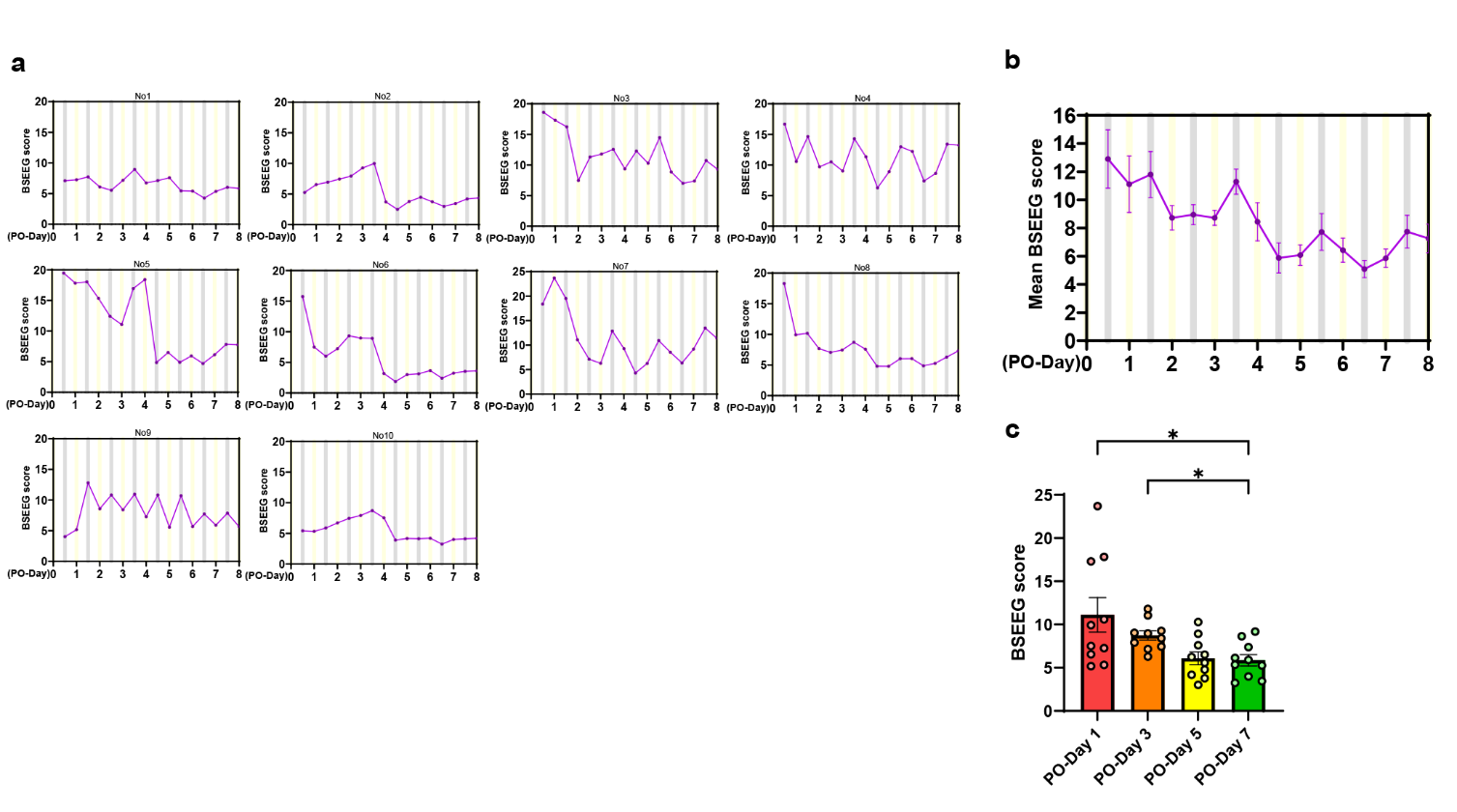

**Fig. S3** The postoperative raw BSEEG score in aged mice. (a) The time courses of postoperative raw BSEEG score in each aged mouse. (b) The time course of mean postoperative raw BSEEG score in aged mice. (c) The differences in raw BSEEG scores comparing PO-Day 1, PO-Day 3, PO-Day 5, and PO-Day 7 groups in aged mice(mean = 11.12 (PO-Day 1), 8.73 (PO-Day 3), 6.085 (PO-Day 5), 5.86 (PO-Day 7); p = 0.023 (PO-Day 1 vs. PO-Day 7), p = 0.021 (PO-Day 3 vs. PO-Day 7))

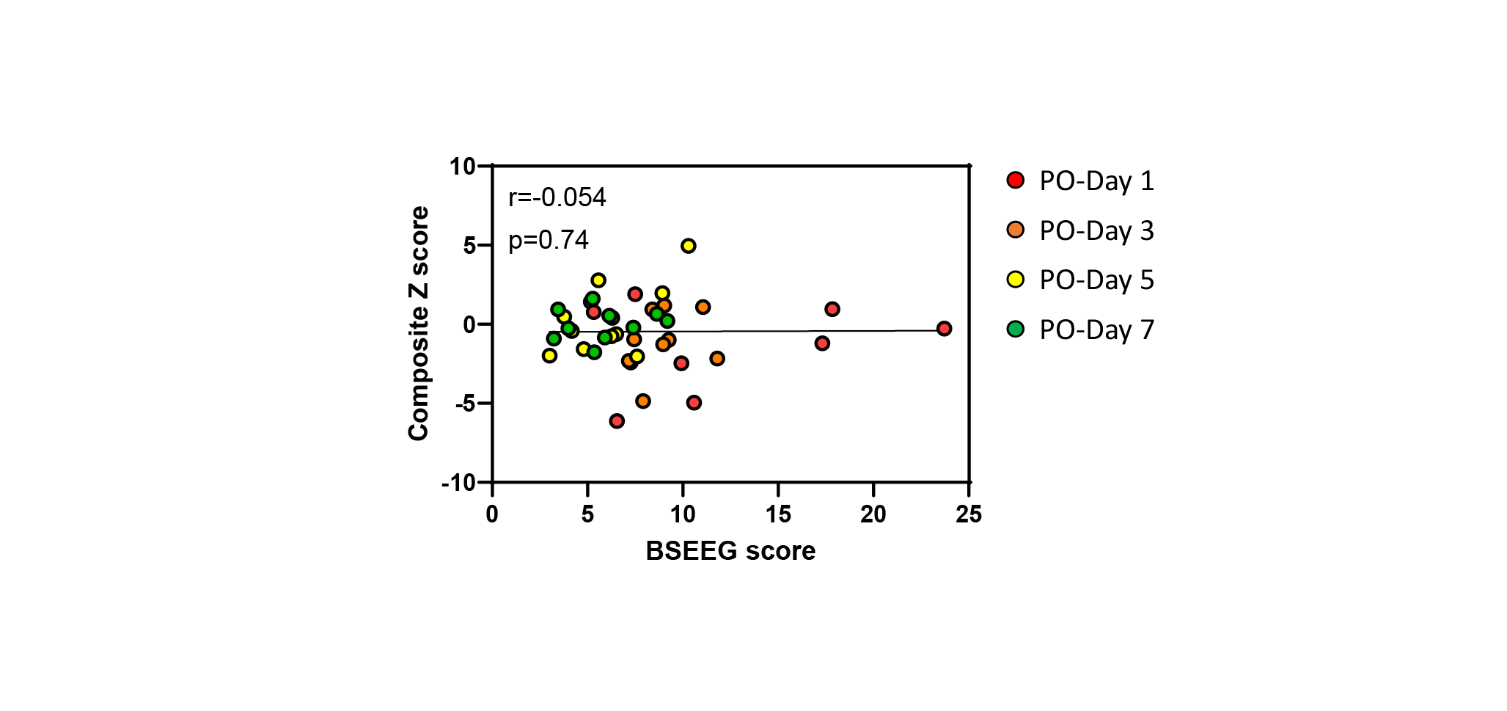

**Fig. S4** Correlation between BSEEG scores and composite Z scores in aged mice
